# A rare variant on a common risk haplotype of *HFE* causes increased risk of hereditary hemochromatosis

**DOI:** 10.1101/547471

**Authors:** Andrew M. Glazer, Lisa Bastarache, Lynn Hall, Laura Short, Tiffany Shields, Brett M. Kroncke, Joshua C. Denny, Dan M. Roden

## Abstract

Hereditary hemochromatosis (HH) is an autosomal recessive disorder of excess iron absorption. The most common form, HH1, is caused by loss of function variants in *HFE. HFE* encodes a cell surface protein that binds to the Transferrin Receptor (TfR1), reducing TfR1’s affinity for the transferrin/iron complex and thereby limiting cellular iron uptake. Two common missense alleles for HH1 have been identified, *HFE* C282Y and *HFE* H63D; H63D is considered to be a less penetrant allele. When we deployed Phenotype Risk Scores (PheRS), a method that aggregates multiple symptoms together in Electronic Health Records (EHRs), we identified *HFE* E168Q as a novel variant associated with HH. E168Q is on the same haplotype as H63D, and in a crystal structure *HFE* E168 lies at the interface of the HFE-TfR1 interaction and makes multiple salt bridge connections with TfR1. In *in vitro* cell surface abundance experiments, the *HFE* E168Q+H63D double mutation surprisingly increased cell surface abundance of HFE by 10-fold compared to wildtype. In coimmunoprecipitation experiments, however, HFE C282Y, E168Q, and E168Q+H63D completely abolished the interaction between HFE and TfR1, while H63D alone only partially reduced binding. These findings provide mechanistic insight to validate the PheRS result that *HFE* E168Q is an HH1-associated allele and lead to the reclassification of E168Q from a variant of uncertain significance to a pathogenic variant, according to ACMG guidelines. *HFE* E168Q results in loss of HFE function by disrupting the HFE-TfR1 interaction. In addition, some disease manifestations attributed to H63D may reflect the functional effects of E168Q.

## Introduction

Hereditary hemochromatosis (HH) is a genetic disorder in which the body absorbs excess iron.^1^ The excess iron can cause damage to the body’s tissues, resulting in a myriad of symptoms, sometimes culminating in liver failure, heart failure, or diabetes. HH has been linked to variants in multiple genes, including autosomal recessively inherited variants in *HFE* (Hemochromatosis Type 1, HH1 [MIM 235200]). Two classic common risk alleles for HH1 in *HFE* are C282Y and H63D.^2,3^ These missense variants are common, with minor allele frequencies of 0-6% and 2-15% respectively, depending on ancestry.^4^ C282Y homozygotes are considered to have the highest risk for HH, with C282Y heterozygotes, H63D homozygotes, and C282Y/H63D compound heterozygotes at lower risk.^5,6^

HFE is a 343 amino acid single-pass transmembrane protein with a large extracellular domain. The extracellular domain of HFE binds the extracellular domain of Transferrin Receptor 1 (TfR1), reducing the affinity of TfR1 for transferrin. When *HFE* is unable to bind TfR1 sufficiently, TfR1 has increased affinity for transferrin, resulting in excess iron uptake into cells and hemochromatosis. C282Y eliminates a disulfide bond in the extracellular domain of HFE, causing misfolding of HFE and failure to traffic to the cell surface.^3^ H63D is thought to cause HH by having a smaller effect on reducing the affinity of TfR1 for transferrin.^2^ Several additional rare variants have been observed in patients with HH in the literature, typically as compound heterozygotes with C282Y or H63D.^7-9^

A promising method to evaluate the risk of genetic variants is in environments that have relatively unselected populations with available Electronic Health Record (EHR) data and genotypic data. Recently, we have developed Phenotype Risk Scores (PheRS) as a method to analyze syndromic phenotypes that have a range of phenotypic effects and to link novel variants. The first deployment of PheRS in a biobank population identified 18 associations between SNPs and syndromic diseases.^10^ One of the associations was between *HFE* E168Q (c.502G>C) and hemochromatosis risk score. Out of the 40 heterozygous carriers of *HFE* E168Q, 8 had highly elevated PheRS’s for HH, and 4 had received a liver transplant. However, whether E168Q actually alters HFE function remains unclear. Largely due to the lack of *in vitro* functional data about its mechanism, E168Q is still classified as a variant of uncertain significance.^10^

Here we investigate the function of *HFE* E168Q using *in vitro* functional assays. We find that E168Q is located on the same haplotype as H63D, and that E168Q+H63D has increased abundance at the cell surface. E168Q lies at the interface of the HFE-TfR1 interaction and completely disrupts that interaction, establishing a mechanism for E168Q’s association with hemochromatosis.

## Material and Methods

### Haplotype analysis

52,573 adult (>18 years) individuals of European ancestry were included in the analysis. These individuals were genotyped with the Multi-Ethnic Global Array (Illumina) or the Infinium HumanExome BeadChip array (Illumina). Genotypes were determined for *HFE* H63D (rs1799945) or *HFE* E168Q (rs146519482). Ancestry was determined from STRUCTURE.^11^

### Mutagenesis and Transfection

pCB6-HFE-EGFP was a gift from Pamela Bjorkman (Addgene plasmid # 12104). This plasmid was mutated with a Quikchange Lightning Multi-site kit (Agilent) to create C282Y, H63D, E168Q, and E168Q+H63D mutant plasmids. Plasmids were transfected into HeLa, HepG2, Chinese Hamster Ovary, or HEK293 cells using Fugene 6 (Promega) following manufacturer’s instructions and studied 48-72 hours post-transfection.

### Cell surface abundance assays

To stain cells for confocal and flow cytometry experiments, HeLa, HepG2, HEK293, or CHO cells were stained with an anti-HFE antibody while still alive—thus, having an intact cell membrane—to quantify HFE at the cell surface. Briefly, cells transfected with wildtype or mutant *HFE*-GFP (see above) were trypsinized, resuspended in complete media, washed in PBS+1%Bovine Serum Albumin, incubated with a polyclonal anti-HFE antibody at 1:500 dilution (ThermoFisher PA5-37364), washed twice in PBS+BSA, incubated with a Alexa Fluor 647-anti-rabbit secondary antibody at 1:500 (ThermoFisher A-21245), and washed twice in PBS+BSA. For flow cytometry, cells were analyzed on a BD LSR Fortessa instrument, using a 488 nm laser and 525/50 nm filter for GFP, and a 633 nm laser and 660/20 filter for Alexa Fluor 647. Single cells were identified from side and forward scatter parameters, and GFP and Alexa Fluor 647 laser levels were set so that untransfected cells had a median of 100 for each. Cells with a high level of GFP were identified (cells with ∼100-fold GFP levels relative to wildtype; 10^1.8 to 10^2.2-fold higher). The median Alexa Fluor 647 level of highly-GFP+ cells was calculated and averaged across at least 3 replicate samples. Statistical analyses were performed in R. Student’s two tailed t-tests were used for comparisons between groups. For confocal microscopy, cells were stained as above, fixed with 4% paraformaldehyde, washed with PBS+Hoechst, and imaged on an Olympus FV-1000 confocal microscope using identical settings for each mutation.

### Co-Immunoprecipitation

HeLa cells were chosen for coimmunoprecipitation experiments because wildtype HFE trafficked best to the cell surface in HeLa cells (Figure S1) and they had been previously used for coimmunoprecipitation experiments.^9^ HeLa cells were cotransfected using Fugene 6 with wildtype or mutant *HFE*-GFP plasmids (see above) and a Transferrin Receptor 1 expression plasmid pcDNA3.2/DEST/hTfR-HA, a gift from Robin Shaw (Addgene plasmid #69610). A Pierce Classic Magnetic IP/Co-IP Kit (ThermoFisher #88804) was used to harvest and coimmunoprecipitate the cells, following manufacturer’s instructions. Cells were precipitated with a rabbit anti-GFP antibody (AbCam #ab290), then Western blots were performed using the anti-GFP antibody (1:2500) or a mouse anti-TfR1 antibody (ThermoFisher #13-6800, 1:500) or a secondary anti-rabbit HRP (Promega #W4011) and anti-mouse HRP (Promega W-4021) antibody, each at 1:10,000.

### Variant classification

*HFE* E168Q was classified according to American College of Medical Genetics and Genomics criteria,^12^ which integrates multiple variables into a benign/likely benign/uncertain significance/likely pathogenic/pathogenic classification. The University of Maryland Genetic Variant Interpretation Tool was used to implement these criteria (medschool.umaryland.edu/Genetic_Variant_Interpretation_Tool1.html).^13^

## Results

### *HFE* E168Q is a rare allele on the same haplotype as H63D

*HFE* E168Q was previously shown using PheRS to have an association with hemochromatosis.^10^ When we examined *HFE* genotypes of E168Q heterozygotes, we observed that E168Q cosegregated with the common H63D allele (88/88 E168Q heterozygotes had at least 1 H63D allele; Table 1). In individuals with European ancestry, E168Q had a minor allele frequency of 0.00084 and H63D had a minor allele frequency of 0.149. E168Q was exclusively present in individuals of European ancestry, except for 1 individual whose ancestry was undetermined by STRUCTURE, and is likely of mixed ancestry. 13 of the 88 E168Q heterozygotes (14.7%) were homozygous for H63D, similar to the minor allele frequency of H63D, indicating that these individuals were likely E168Q+H63D / H63D compound heterozygotes. Together, these data indicate that E168Q is a rare variant that arose on the H63D allele of *HFE*.

**Table 1:** E168Q is a rare variant on the same haplotype as H63D. Co-occurrence of *HFE* H63D and E168Q genotypes in 52,573 adult European-ancestry individuals. The 168Q allele only appears in the presence of the 63D allele, indicating that the 168Q allele is on the same haplotype as the 63D allele.

|  | H63D allele |  |  |
| --- | --- | --- | --- |
| E168Q allele | HH | HD | DD |
| EE | 38,084 | 13,256 | 1,145 |
| EQ | 0 | 75 | 13 |
| QQ | 0 | 0 | 0 |

### *HFE* E168Q+H63D has increased abundance at the cell surface

A common mechanism for loss of function of membrane proteins is a defect in trafficking to the cell surface, often due to misfolding of the protein and subsequent aggregation along the secretory pathway.^14^ Cell surface abundance can also be affected by altered rates of internalization or degradation. We developed a dual flow cytometry and confocal microscopy-based assay to assay the subcellular localization of a GFP-tagged HFE protein. Testing of four cell lines (HeLa, HepG2, Chinese Hamster Ovary, and HEK293) revealed that wildtype HFE trafficking efficiency varied widely between cell lines, trafficking best in HeLa cells, followed by HepG2 cells (Figure S1). To test whether mutant HFE proteins were present in different abundances at the cell surface, we examined the cellular localization of GFP-tagged HFE wildtype protein and HFE mutants H63D, E168Q, C282Y, and the E168Q+H63D double mutant (hereafter referred to as E168Q+H63D) (Figure 1). C282Y showed a dramatic trafficking defect (5% of wildtype level, p=3.2e-5, two-tailed T test). E168Q had a mild but significant increase in surface abundance (136% of wildtype level, p=0.01). H63D surprisingly had a higher abundance than wildtype (722% of wildtype level, p=0.02). E168Q+H63D also had a large increase in surface abundance (970% of wildtype level, p=0.02). Similar relative surface abundance results were observed in the HepG2 liver cell line, albeit with lower overall levels of surface trafficking (Figures 1, S1). Therefore, in contrast to C282Y, H63D and H63D+E168Q have increased cell surface abundances.

**Figure 1:**
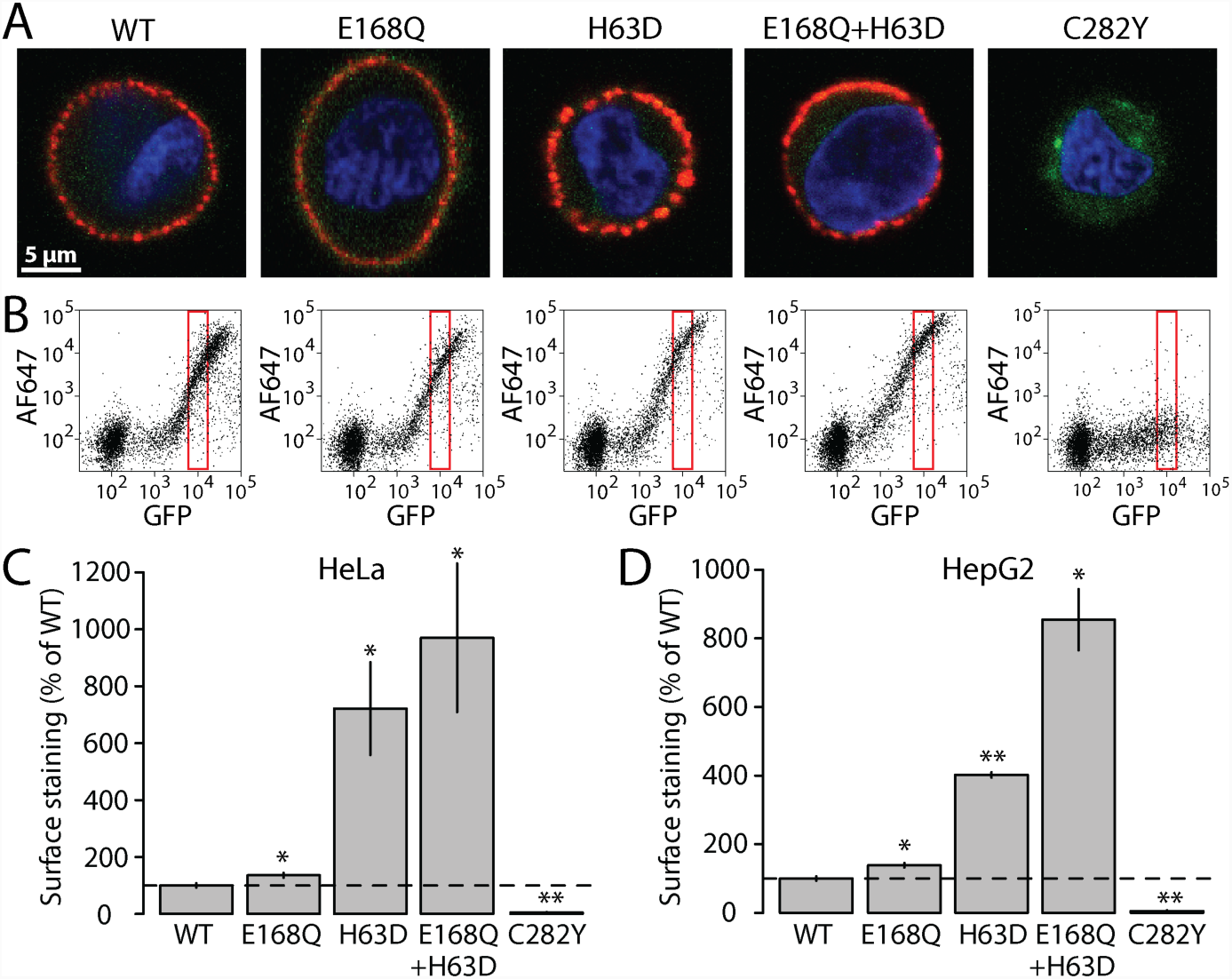
HFE E168Q+H63D has enhanced trafficking to the cell surface. HeLa or HepG2 cells were transfected with *HFE*-GFP and stained while alive for HFE at the cell surface (Alexa Fluor 647). A) Representative confocal microscopic images of HeLa cells (blue=Hoechst nuclear stain, green=GFP, red=Alexa Fluor 647). B) Representative flow cytometry plots of surface HFE staining in HeLa cells. Red box indicates transfected cells (GFP+ 100-fold higher than untransfected cells). X and Y-axis scales are in log10 units. C-D) Quantification of flow cytometry data of cell surface staining in HeLa (C) or HepG2 (D) cells. Values are normalized to the mean wildtype level. *p<0.05, **p<0.005, two-tailed T test. C) n=5-6 replicate samples/mutation. D) n=3 replicate samples/mutation.

### HFE E168 is at the interface of the HFE-Transferrin Receptor interaction

We next examined the location of E168Q within the HFE protein. A crystal structure of the HFE-TfR1 interaction has been solved, together with the HFE binding partner B2M (Figure 2).^15^ That crystal structure revealed six residues of HFE that made salt bridges with TfR1 (Table S1). Intriguingly, in the structure, one of these residues is HFE E168, which is located at the HFE-TfR1 interface, and makes salt bridges with two TfR1 residues (TfR1-R629 and TfR1-Q640). HFE E168Q is also predicted to make an intramolecular salt bridge with HFE N108.

**Figure 2:**
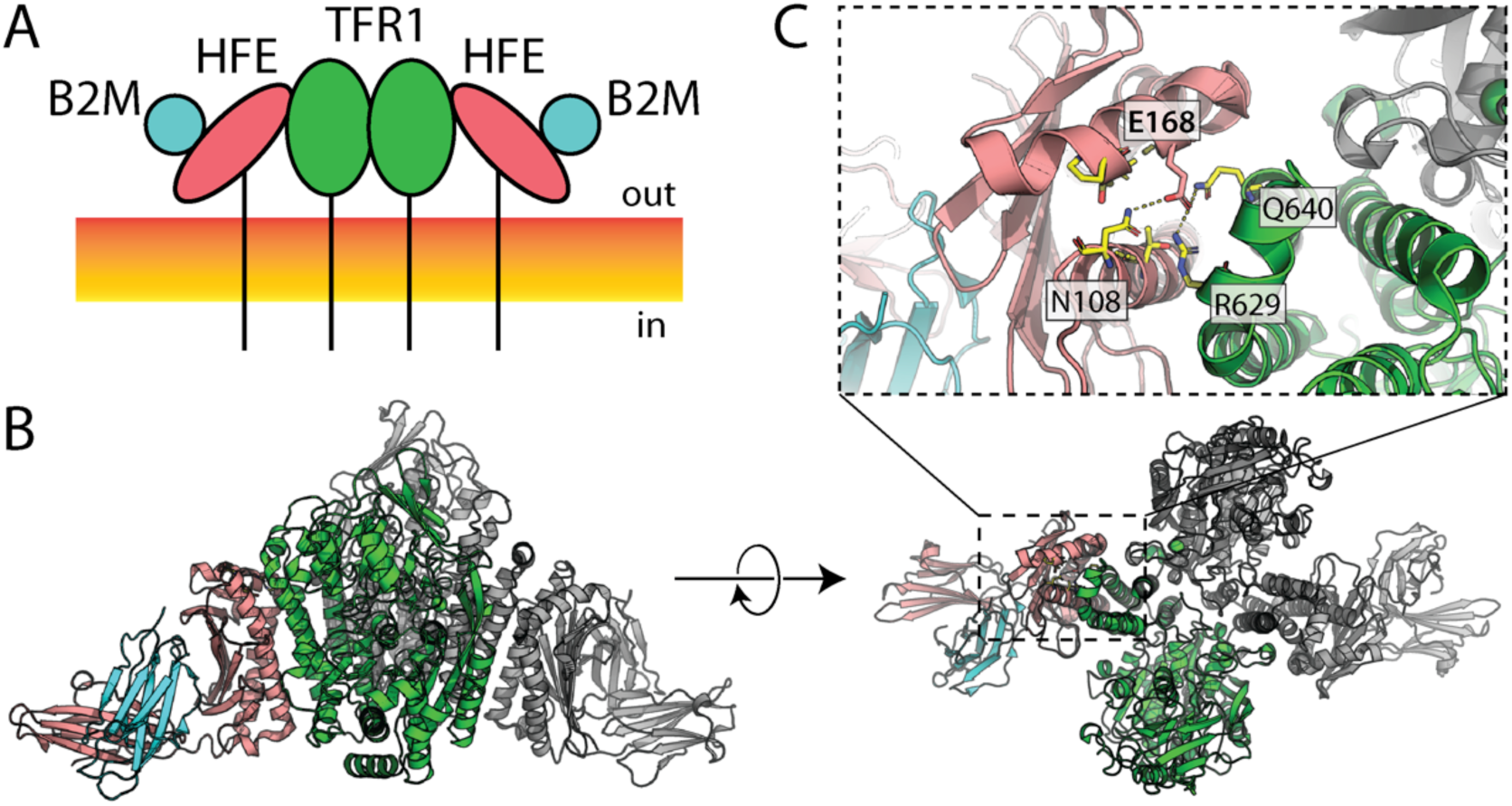
HFE E168 is located at the interface of the HFE-TfR1 interaction. A) Schematic of HFE/TfR1/B2M interactions at the cell surface. Orange/yellow rectangle indicates plasma membrane. B-D) Crystal structure of HFE-TfR1-B2M extracellular domains.^15^ HFE is colored in salmon, TfR1 in green, B2M in blue. Each protein is present twice in the complex, but only one instance is colored, and the other instance is shown in gray. B) Side view. C) Top view with zoom. Salt bridges between HFE E168 and HFE N108, TfR1 R629, and TfR1 Q640 are shown with dotted lines.

### HFE E168Q disrupts the interaction between HFE and the Transferrin Receptor

Because of HFE E168’s location and contacts with TfR1, we hypothesized that the HFE E168Q variant disrupts the binding between HFE and TfR1. To test this, we performed coimmunoprecipitation experiments, precipitating HFE-GFP using an anti-GFP antibody and measuring coimmunoprecipitation of TfR1 (Figure 3). Wildtype HFE-GFP coimmunoprecipitated TfR1, but C282Y, E168Q, and E168Q+H63D showed no coimmunoprecipitation of TfR1. Across multiple replicates, C282Y had lower overall intensity of anti-GFP staining in both input and immunoprecipitated samples, consistent with a previously observed accelerated degradation rate of this variant.^3^ H63D showed a detectable but decreased coimmunoprecipitation of TfR1. Thus, E168Q and E168Q+H63D had a more severe defect in binding TfR1 than H63D.

**Figure 3:**
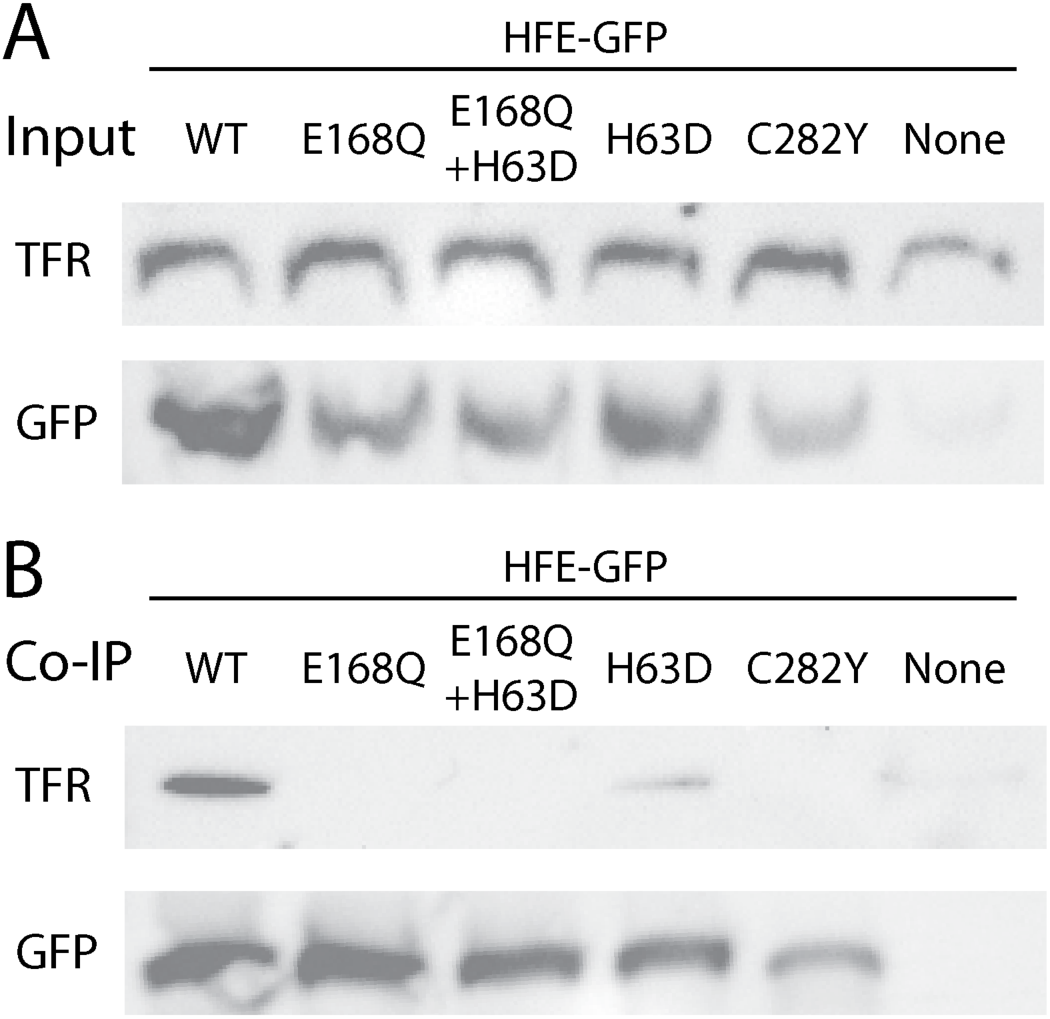
HFE E168Q disrupts the interaction between HFE and the Transferrin Receptor. Coimmunoprecipitation experiments. A) Input. B) HFE-GFP was immunoprecipitated with an anti-GFP antibody. For A and B, Western blots using anti-GFP or anti-TfR1 are shown. Similar results were observed across 3 replicate experiments.

### Pathogenicity reclassification of *HFE* E168Q

We used American College of Medical Genetics and Genomics criteria to determine the classification of *HFE* E168Q using data available before this study and after this study (Table 2). Despite the genetic association in a biobank population, *HFE* E168Q before this study was still classified as a variant of uncertain significance based on criterion PS4 (variant prevalence in affected individuals is significantly increased compared with the prevalence in controls).^10^ Based on the updated *in vitro* functional data in this paper (criteria PS3), *HFE* E168Q now has enough evidence to be classified as a pathogenic variant.

**Table 2.** Classification of HFE E168Q using ACMG guidelines. American College of Medical Genetics and Genomics (ACMG) classification guidelines^12^ were implemented using the online Genetic Variant Interpretation Tool.^13^ Because of the *in vitro* functional evidence in this study, the classification of E168Q changed from Variant of Uncertain Significance (VUS) to Pathogenic.

| ACMG<br>Criterion | Description | Before<br>this study | After this<br>study |
| --- | --- | --- | --- |
| PS3 | Well-established *in vitro* or *in vivo* functional studies supportive of a damaging effect on the gene or gene product | No | Yes |
| PS4 | Variant prevalence in affected individuals is significantly increased compared with the prevalence in controls | Yes | Yes |
| Classification |  | VUS | Pathogenic |

## Discussion

### Phenotype Risk Scores for syndromic traits

*HFE* E168Q was identified as a novel HH1 allele with PheRS, a recently developed method that combines multiple phenotypes into a weighted score, to study the HH risk of different *HFE* variants. The PheRS for hemochromatosis includes 22 symptoms, such as liver cirrhosis, hepatic cancer, cardiac dysrhythmias, and type 2 diabetes. The *in vitro* work presented here validating E168Q as a loss of function allele validates the use of PheRS as a powerful way to assess the disease risk of variants. As the number of individuals in EHR datasets linked to genotyping grows, this approach will gain in power to detect genetic associations. Given the size of contemporary biobanks linking DNA variation to human phenotypes,^16^ this approach will likely prove fruitful for rare but not ultrarare variants such as E168Q (minor allele frequency of 0.08% in Europeans) that are still present in many individuals in large biobanks. However, for some variants like *HFE* E168Q, statistical association in biobanks is not enough to classify variants as pathogenic or likely pathogenic, and further *in vitro* functional validation is required. The combination of statistical association by PheRS and *in vitro* loss of function phenotype is enough to reclassify *HFE* E168Q as pathogenic.

### Mechanism of *HFE* E168Q

A main mechanism of transmembrane proteins having loss of function is misfolding and subsequent failure to traffic to the cell membrane. However, in this work, we surprisingly observed that HFE E168Q+H63D had increased abundance at the cell surface. Much of this surface abundance difference was due to H63D, although E168Q alone had a mild but significant increase in surface abundance. We observed that HFE E168 was at the interface of the HFE-TfR1 interaction and made multiple salt bridge contacts with TfR1, suggesting that E168Q would disrupt the salt bridge contacts with TfR1. Indeed, coimmunoprecipitation experiments showed a complete loss of binding of E168Q and E168Q+H63D. Although it is difficult to predict the exact configuration of the mutant glutamine in the crystal structure, the glutamine likely completely disrupts the interaction between HFE-168 and TfR1-R629 and likely alters the contacts with TfR1-Q640. Two main alpha helices make contact with TfR1, termed α1 and α2.^15^ HFE E168 is located in α2. Previous work showed that mutation to alanine of two residues in α1, V100 and W103 (called V78 and W81 in the original paper), also abrogated the binding between HFE and TfR1.^17^ Therefore, we propose a model in which *HFE* E168Q is unable to bind TfR1 and TfR1 therefore has an increased affinity for transferrin, causing iron overload and HH1.

### Improved prediction of HH risk

Our results suggest a template for improved prediction of HH risk. Our results further suggest that genotyping for H63D and C282Y alone might not be sufficient to determine HH1 risk. H63D is considered to be a low/variable penetrance HH1 allele,^5,6^ and E168Q presence may underlie some of the HH1 risk previously attributed to H63D alone and explain some of its variable penetrance.^18^ Other rare HFE variants in α1 and α2 or making salt bridge connections with TfR1 may also disrupt TfR1 binding and lead to hemochromatosis. Integrated phenotyping methods like PheRS show promise to identify risk variants and may also identify patients with underrecognized disease. However, further functional studies are often necessary to validate these variants; 9/18 variants in the initial PheRS paper were classified as variants of uncertain significance despite their statistical association with disease.^10^ *In vitro* functional studies such as the surface abundance and coimmunoprecipitation studies in this paper can validate the genetic results and result in reclassification of variants as benign or pathogenic. We anticipate that these methods will be more broadly applied to other variants, genes, and diseases to better predict disease risk.

## Supplemental Data Description

The Supplemental Data contains 1 figure and 1 table.

**Figure S1.**
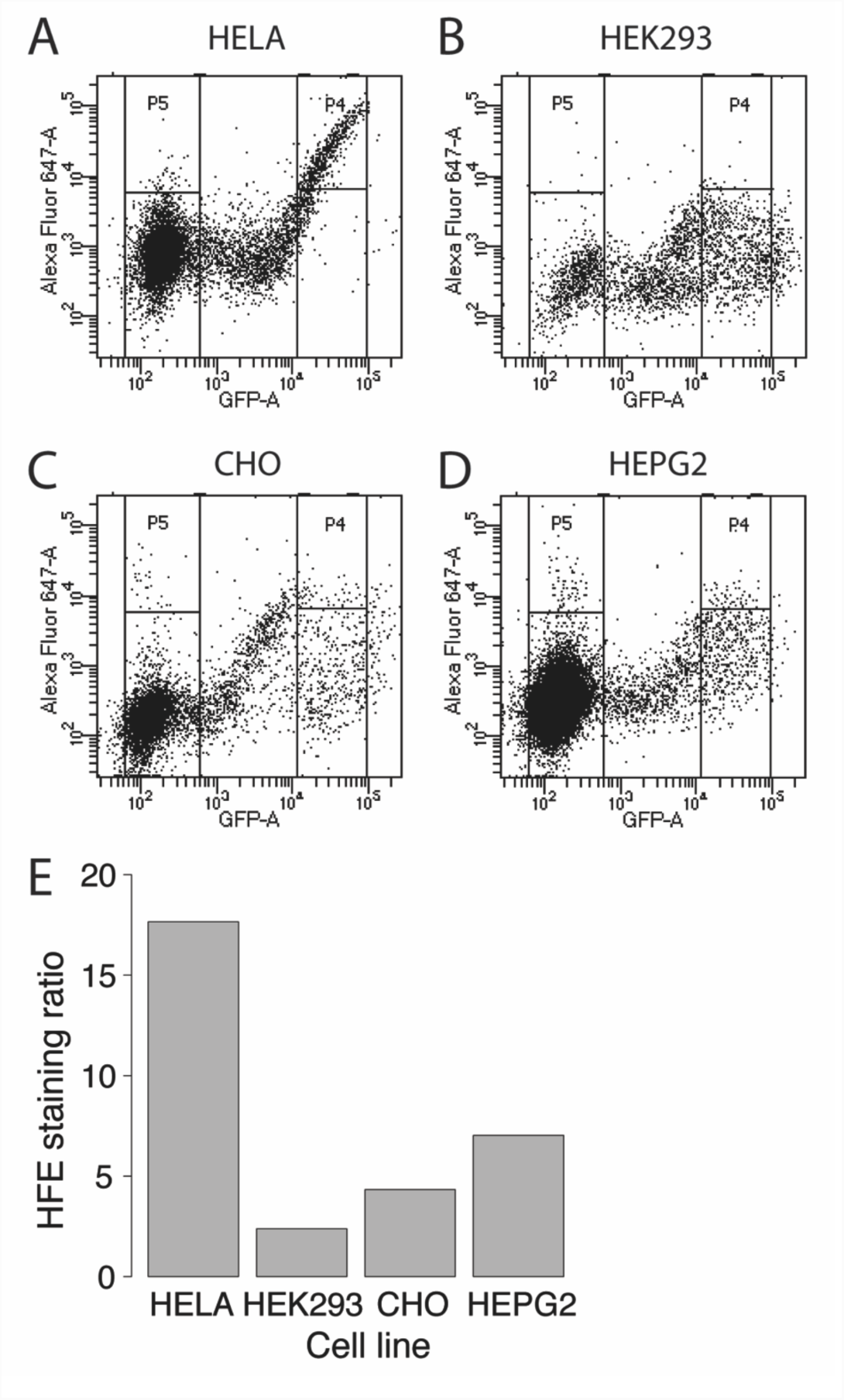
HFE traffics robustly to the cell membrane in HeLa cells. A-D) Wildtype HFE cell surface staining (Alexa Fluor 647) vs. HFE-GFP level (GFP) in transiently transfected HeLa (A), HEK293 (B), CHO (C), or HepG2 cells (D). E) A HFE staining ratio was calculated by taking the median Alexa Fluor 647 level in highly expressing cells (P4) divided by the median Alexa Fluor 647 level in untransfected cells (P5).

**Table S1.** Residues of HFE forming salt bridges with TfR1.

| **HFE residue**<br>**(contemporary**<br>**numbering)** | **HFE residue**<br>**(Bennett *et al.***<br>**numbering)** | **HFE location** | **Interacting TfR1**<br>**residue** |
| --- | --- | --- | --- |
| Gln86 | Gln64 | α1 helix | Thr658 |
| Glu107 | Glu85 | α1 helix | Arg629 |
| Asn108 | Asn86 | α1 helix | Arg629 |
| Glu168 | Glu146 | α2 helix | Arg629, Gln640 |
| Arg175 | Arg153 | α2 helix | Gln640 |
| Gln178 | Gln156 | α2 helix | Asp648 |

Table adapted from Bennett *et al.* 2000.<sup>15</sup>

## Acknowledgements

This work was supported by grants R01-LM010685 from the National Library of Medicine and P50-GM115305 from the National Institute for General Medical Sciences and a grant from the Robert J. Kleberg, Jr. and Helen C. Kleberg Foundation. BioVU received and continues to receive support through the National Center for Research Resources (UL1-RR024975), which is now the National Center for Advancing Translational Sciences (UL1-TR000445). A.M.G. was supported by F32 HL137385 and T32 HG008341. B.M.K. was supported by K99 HL135442.

